# Sex differences in behavioral responding and dopamine release during Pavlovian learning

**DOI:** 10.1101/2021.10.04.463059

**Authors:** Merridee J. Lefner, Mariana I. Dejeux, Matthew J. Wanat

## Abstract

Learning associations between cues and rewards requires the mesolimbic dopamine system. The dopamine response to cues signals differences in reward value in well-trained animals. However, these value-related dopamine responses are absent during early training sessions when cues signal differences in the reward rate. These findings suggest cue-evoked dopamine release conveys differences between outcomes only after extensive training, though it is unclear if this is unique to when cues signal differences in reward rate, or if this is also evident when cues signal differences in other value-related parameters such as reward size. To address this, we utilized a Pavlovian conditioning task in which one audio cue was associated with a small reward (one pellet) and another audio cue was associated with a large reward (three pellets). We performed fast-scan cyclic voltammetry to record changes in dopamine release in the nucleus accumbens of male and female rats throughout learning. While female rats exhibited higher levels of conditioned responding, a faster latency to respond, and elevated post-reward head entries relative to male rats, there were no sex differences in the dopamine response to cues. Multiple training sessions were required before cue-evoked dopamine release signaled differences in reward size. Reward-evoked dopamine release scaled with reward size, though females displayed lower reward-evoked dopamine responses relative to males. Conditioned responding related to the decrease in the peak reward-evoked dopamine response and not to cue-evoked dopamine release. Collectively these data illustrate sex differences in behavioral responding as well as in reward-evoked dopamine release during Pavlovian learning.

## Introduction

Efficient reward seeking involves identifying cues that predict rewards and discriminating between cues that signal different reward options. The mesolimbic dopamine system plays an integral role in regulating behavioral responses toward reward-associated cues [1,2]. Cue-evoked dopamine responses convey reward-related information such as the relative reward size [3-5], reward probability [6,7], and reward rate [8]. While this effect is evident in extensively trained animals, the emergence of these signals during initial training sessions has not been well-characterized. We recently utilized a Pavlovian conditioning task to demonstrate that cue-evoked dopamine release encodes reward rate (i.e., the time elapsed since the previous reward delivery) after extensive training (>24 sessions) [8], but not during the first six training sessions [9]. These findings suggest cue-evoked dopamine encodes reward rate through a multistep process: by first signaling an upcoming reward independent of value during initial training sessions, and after additional training conveying the relative difference in value between cues. However, it remains unclear if extensive training is similarly required for cue-evoked dopamine signals to convey prospective value-related parameters, such as reward size.

The aforementioned research was primarily performed in male subjects, however increasing evidence highlights sex differences in behavioral responding. Across studies female subjects display augmented behavioral responses compared to males, including higher locomotor activity, faster latency, and elevated conditioned responding [10-20]. Furthermore, prior research has identified sex differences in dopamine neuron activity and release [21-26]. These differences in dopamine transmission between males and females could account for the observed sex differences in dopamine-dependent behaviors [13-20,27]. However it is not known if sex differences during Pavlovian learning are accompanied by distinct patterns of dopamine signaling.

In this study, we trained male and female rats on a Pavlovian task where one cue was associated with a small reward (one pellet) and another cue was associated with a large reward (three pellets). Female rats displayed higher levels of conditioned responding, a faster latency to the food port, and a higher number of post-reward head entries compared to male rats. We used fast-scan cyclic voltammetry to record changes in dopamine release in the nucleus accumbens (NAc) throughout learning. The cue-evoked dopamine response did not encode differences in reward size during the first six training sessions, but did signal differences in value during later sessions. There were no differences in cue-evoked dopamine release between males and females. In contrast, the dynamics of reward-evoked dopamine release was influenced by both reward size and sex. Both male and female rats displayed higher reward-evoked dopamine release to the larger reward option, though females exhibited lower reward-evoked dopamine levels compared to males. These data illustrate that sex differences in dopamine transmission are stimulus-specific.

## Methods

### Subjects and surgery

All procedures were approved by the Institutional Animal Care and Use Committee at the University of Texas at San Antonio. Male (300-350 g) and female (200-250 g) Sprague-Dawley rats (Charles River, MA) were pair-housed upon arrival and given ad libitum access to water and chow and maintained on a 12-hour light/dark cycle (n = 8 male rats/9 electrodes, 5 female rats/5 electrodes). Carbon fiber voltammetry electrodes consisted of a carbon fiber housed in silica tubing and cut to a length of ∼150 μm [28]. Voltammetry electrodes were surgically implanted to target the NAc (relative to bregma: 1.3 mm anterior; ± 1.3 mm lateral; 7.0 mm ventral) along with a Ag/AgCl reference electrode. Rats were single-housed following surgery and allowed to recover for >3 weeks before beginning training.

### Behavioral procedures

After recovering from surgery, rats were placed and maintained on mild food restriction (∼15 g/day of standard lab chow) to target 90% free-feeding weight, allowing for an increase of 1.5% per week. Behavioral sessions were performed in chambers (Med Associates) that had grid floors, a house light, a food tray, and auditory stimulus generators (2.5 and 4.5 kHz tones). To familiarize rats with the chamber and food retrieval, rats underwent a single magazine training session in which 20 food pellets (45 mg, BioServ) were non-contingently delivered at a 90 ± 15 s variable interval. Rats then underwent up to 9 Pavlovian conditioning sessions (1/day) that each consisted of 50 trials where the termination of a 5 s audio cue (CS; 2.5 kHz tone or 4.5 kHz tone, counterbalanced across animals) resulted in the delivery of a single food pellet (Small Reward trials) or three food pellets (Large Reward trials) and illumination of the food port light for 4.5 s. The three food pellets on Large Reward trials were delivered within 0.4 s after the end of the CS presentation. Each session contained 25 Small Reward trials and 25 Large Reward trials delivered in a pseudorandom order, with a 45 ± 5 s inter-trial interval between all trials. Conditioned responding was quantified as the change in the rate of head entries during the 5 s CS relative to the 5 s preceding the CS delivery [8,9]. We also quantified the latency to initiate a head entry during the CS. For the post-US analysis we calculated the average number of head entries made during a 9 s post-US delivery time window. For more detailed analysis, we also broke the 9 s time window into two 4.5 s epochs that corresponded to when the food tray light was illuminated (Early US; 0-4.5 s) and an equivalent period of time when the food tray light was turned off (Late US; 4.5-9 s).

### Voltammetry recordings and analysis

Chronically-implanted electrodes were connected to a head-mounted amplifier to monitor changes in dopamine release in behaving rats using fast-scan cyclic voltammetry as described previously [8,9,28-32]. The carbon fiber electrodes were held at −0.4 V (vs. Ag/AgCl) with voltammetric scans applied at 10 Hz in which the potential was ramped in a triangular waveform to +1.3 V and back to −0.4 V at a rate of 400 V/s. A principal component regression analysis [33] was performed on the voltammetry signal using a standard training set that accounts for dopamine, pH, and drift. The average post-implantation sensitivity of electrodes (34 nA/μM) was used to estimate dopamine concentration [28]. Chemical verification of dopamine was achieved by obtaining a high correlation of the cyclic voltammogram during a reward-related event to that of a dopamine standard (correlation coefficient r^2^ ≥ 0.75 by linear regression). Voltammetry data for a session were excluded from analysis if the detected voltammetry signal did not satisfy this chemical verification criteria [8,9]. Voltammetry data for a given trial were excluded if the principal component regression analysis failed to extract dopamine current on > 25% of the data points for a given trial (i.e. the residual Q value for the regression analysis exceeded the 95.5% confidence limit for the training set) [9,29,33,34].

The CS-evoked dopamine response was quantified as the average dopamine response during the 5 s CS relative to the 5 s prior to the CS delivery [8,9]. The peak US-evoked dopamine response was quantified as the maximum dopamine response in the 3 s following US delivery relative to 0.5 s prior to US delivery. The area under the curve (AUC) of the post-US dopamine response was quantified as the average dopamine response in the 9 s following US delivery relative to 0.5 s prior to US delivery. To determine potential differences in the decay of reward-evoked dopamine release between males and females, we normalized and aligned to the peak US dopamine response following the Small Reward delivery of the first session. These data were then fit to a single-phase decay curve to calculate the tau for each electrode [32].

### Data analysis

Statistical analyses were performed in GraphPad Prism 9 and RStudio. Behavioral responding and dopamine quantification were analyzed using a mixed-effects model fit (restricted maximum likelihood method), repeated measures where appropriate, followed by a post hoc Sidak’s test. The Geisser-Greenhouse correction was applied to address unequal variances between groups. A repeated measures correlation was used to correlate dopamine signals and behavioral outcomes across all subjects, trial types, and training sessions [9,35]. The full list of statistical analyses is presented in the **Supplementary Table**.

### Histology

Rats were deeply anesthetized, and electrical lesions were applied to the voltammetry electrodes followed by intracardial perfusion with 4% paraformaldehyde. Brains were removed and postfixed for at least 24 hours, then subsequently placed in 15% and 30% sucrose solutions in phosphate-buffered saline. Brains were then flash frozen on dry ice, coronally sectioned, and stained with cresyl violet. Electrode locations were mapped onto a standardized rat brain atlas.

## Results

Rats were trained on a Pavlovian conditioning task in which one audio cue (CS) signaled the delivery of a single sucrose pellet (US; Small Reward trial) and another audio cue signaled the delivery of three sucrose pellets (Large Reward trial, **Fig. 1A**). Conditioned responding was quantified as the change in the rate of head entries during the 5 s CS relative to the rate of head entries during the 5 s preceding the CS [8,9,29]. Rats increased conditioned responding across sessions, with no difference between Small and Large Reward cues (three-way mixed-effects analysis; session effect: *F*_(2.26, 24.86)_ = 14.01, *p* < 0.0001; reward size effect: *F*_(1, 11)_ = 0.03, *p* = 0.86; *n* = 13 rats, **Fig. 1B**). There was a trend for enhanced conditioned responding in female rats (sex effect: *F*_(1, 55)_ = 3.90, *p* = 0.05; session x sex interaction: *F*_(5, 55)_ = 2.34, *p* = 0.05; **Fig. 1B**).

**Figure 1.**
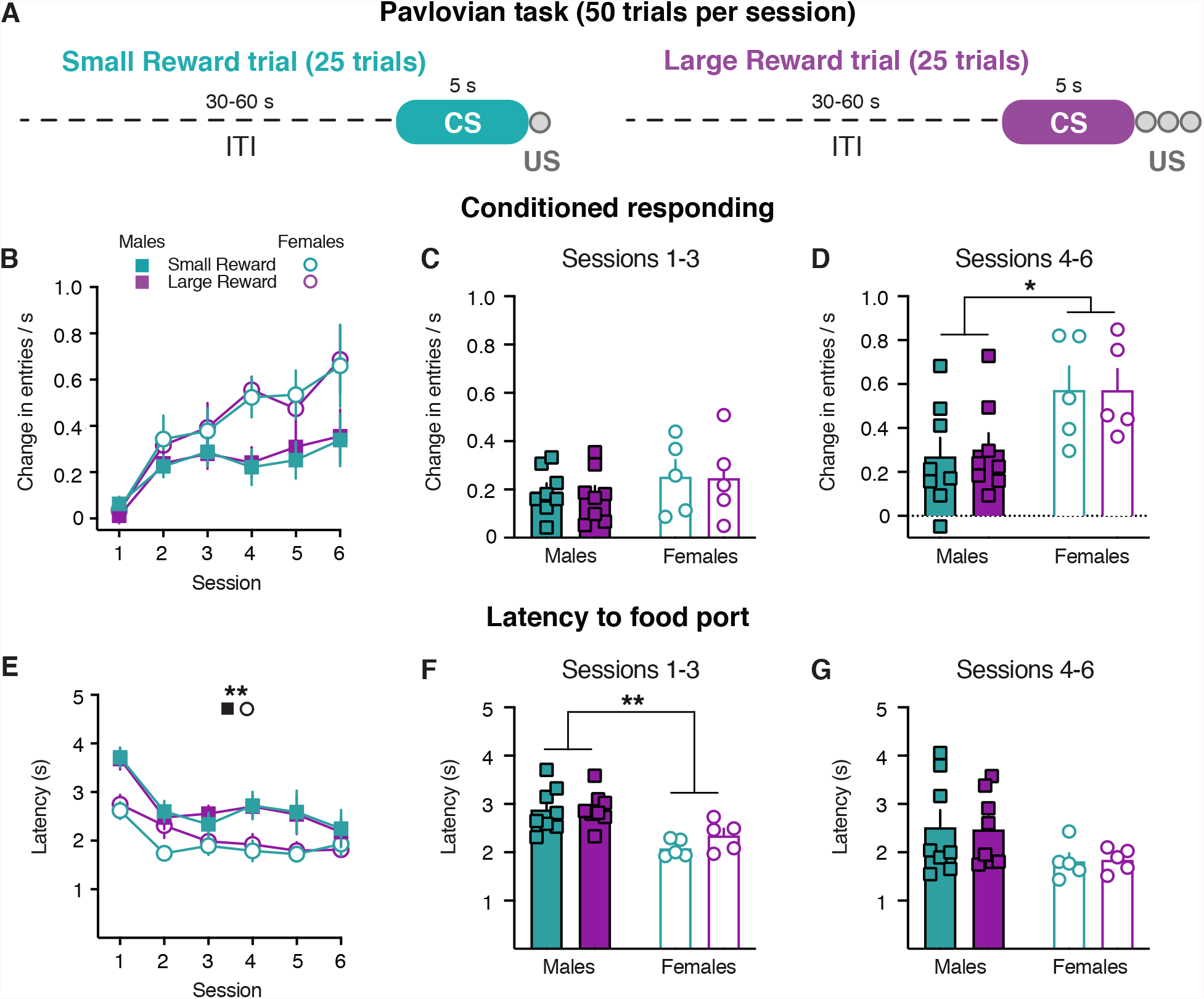
Sex differences in behavioral responding during CS presentation. (A) Training schematic for Pavlovian reward size task. (B) Conditioned responding for males (filled squares) and females (open circles) during Small Reward (teal) and Large Reward (purple) trials. (C) Conditioned responding averaged across the first three sessions of training. (D) Conditioned responding averaged across the latter three sessions of training. (E) Latency to respond to the food port (F) Latency to respond averaged across the first three sessions of training. (D) Latency to respond averaged across the latter three sessions of training. * *p* < 0.05, * * *p* < 0.01.

Rats also decreased the latency to the food port across training sessions, with no difference between Small and Large reward trials (three-way mixed-effects analysis; session effect: *F*_(2.94, 32.33)_ = 6.26, *p* < 0.002; reward size effect: *F*_(1, 11)_ = 0.51, *p* = 0.49; **Fig. 1E**). Females displayed a faster latency to respond across sessions compared to males (sex effect: *F*_(1, 55)_ = 8.80, *p* = 0.004; **Fig. 1E**), consistent with prior findings [19,20]. We further analyzed these behavioral responses when averaged into three-session bins. During the first three sessions there were no sex differences in conditioned responding (two-way mixed-effects analysis; sex effect: *F*_(1, 11)_ = 0.90, *p* = 0.36; **Fig. 1C**), though females exhibited a faster latency to enter the food port (two-way mixed-effects analysis; sex effect: *F*_(1, 11)_ = 14.56, *p* = 0.003; **Fig. 1F**). During the latter three sessions, female rats displayed higher levels of conditioned responding (two-way mixed-effects analysis; sex effect: *F*_(1, 11)_ = 5.11, *p* < 0.05; **Fig. 1D**), though there were no sex differences in the latency to respond (two-way mixed-effects analysis; sex effect: *F*_(1, 11)_ = 2.71, *p* = 0.13; **Fig. 1G**). Collectively these findings illustrate that female rats display augmented behavioral responding within the CS presentation compared to male rats during the first six training sessions of Pavlovian learning. However, these behavioral responses during the cue presentation did not reflect differences in the upcoming reward size.

Given the sex differences in CS-evoked behavior, we next examined if male and female rats differed in their behavioral responses following the reward delivery. Female rats performed a higher number of non-CS head entries relative to males (**Supplementary Fig. 1**), which suggests females performed more head entries following the US. To address this possibility, we examined the head entries performed in the 9 s after the reward was delivered (**Fig. 2A**). Female rats exhibited greater post-US head entries compared to male rats (sex effect: *F*_(1, 55)_ = 17.44, *p* = 0.0001; **Fig. 2B**). Additionally, rats performed more head entries following the delivery of the Large Reward (three-way mixed-effects analysis; reward size effect: *F*_(1, 11)_ = 10.15, *p* = 0.009; **Fig. 2B and Supplementary Fig. 2**). We further examined the post-US head entries in two separate epochs that corresponded to when the food tray light was illuminated (Early US: 0-4.5 s) and an equivalent period of time when the food tray light was turned off (Late US: 4.5-9 s; **Fig. 2A**). During the Early US epoch, female rats made a greater number of head entries compared to male rats (three-way mixed-effects analysis; sex effect: *F*_(1, 55)_ = 13.60, *p* = 0.0005; **Fig. 2C and Supplementary Fig. 2**). During the Late US epoch, rats performed more head entries following the Large Reward delivery (three-way mixed-effects analysis; reward size effect: *F*_(1, 11)_ = 24.65, *p* = 0.0004; **Fig. 2D and Supplementary Fig. 2**). Furthermore, there was a sex x reward size interaction effect as female rats continued to demonstrate a greater number of head entries than males throughout the Late US epoch (sex effect: *F*_(1, 55)_ = 12.20, *p* = 0.001; sex x reward size effect: *F*_(1, 55)_ = 5.94, *p* = 0.02**; Fig. 2D and Supplementary Fig. 2**). Taken together these results illustrate that sex and reward size influences the number of post-US head entries.

**Figure 2.**
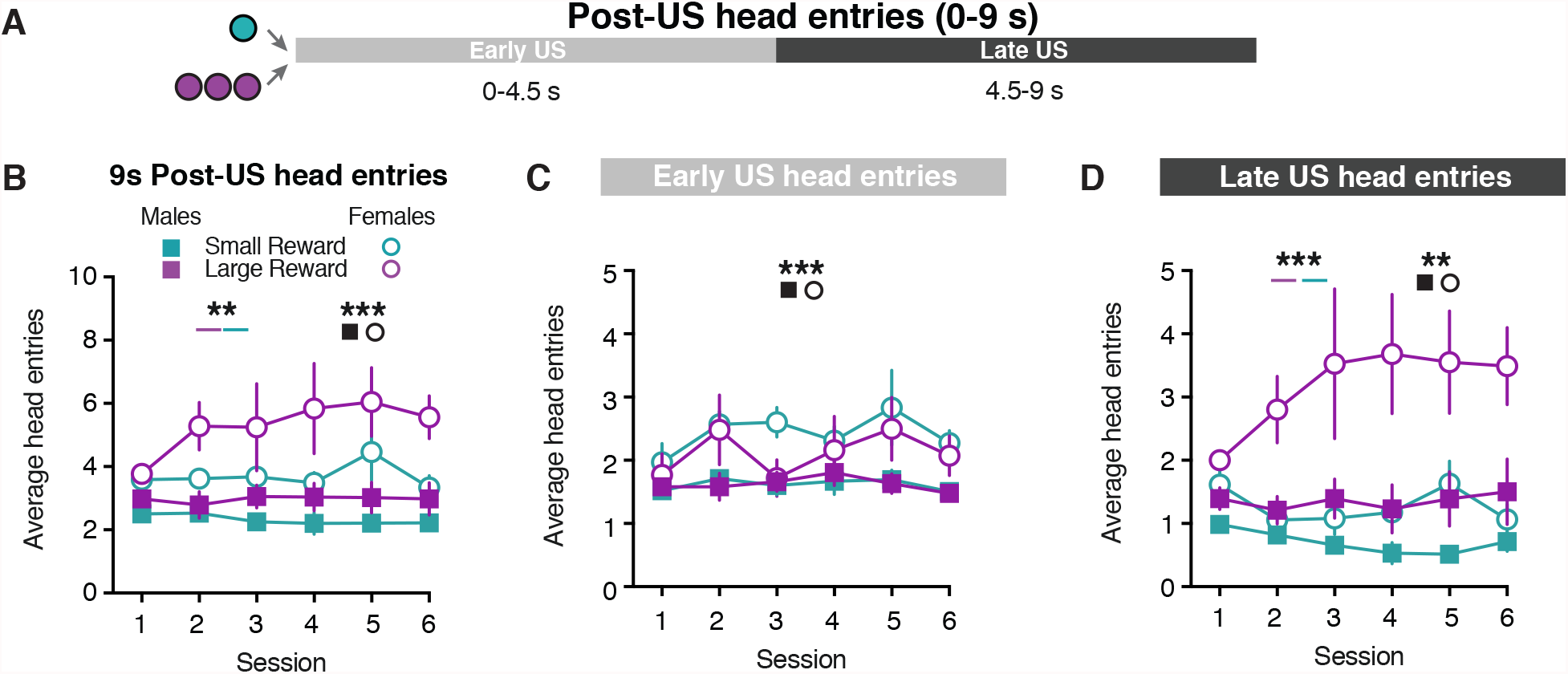
Sex differences in behavioral responding during US presentation. (A) Schematic for post-US epochs: Early US (0-4.5 s) and Late US (4.5-9 s). (B) Average head entries made during the full 9 s post-US window for Small Reward and Large Reward trials. (C) Average head entries made during the Early US for Small Reward and Large Reward trials. (D) Average head entries made during the Late US for Small Reward and Large Reward trials. * * *p* < 0.01, * * * *p* < 0.001

The emergence of Pavlovian conditioned responses depends upon dopamine signaling within the ventral striatum [36]. Here, we performed voltammetry recordings in the NAc to examine how the CS- and US-evoked dopamine responses progressed across training (**Fig. 3A-B**). Both male and female subjects exhibited dopamine release to the CS presentation (**Fig. 3C**). We quantified CS-evoked dopamine release as the average response during the 5 s CS relative to the 5 s prior to the CS, identical to the manner in which conditioned responding was calculated (**Fig. 1**). CS-evoked dopamine release did not differ between sexes or trial type (three-way mixed-effects analysis; session effect: *F*_(1.97, 23.64)_ = 3.22, *p =* 0.06; sex effect: *F*_(1, 30)_ = 0.07, *p* = 0.80; reward size effect: *F*_(1, 12)_ = 3.54, *p* = 0.09; **Fig. 3D**).

**Figure 3.**
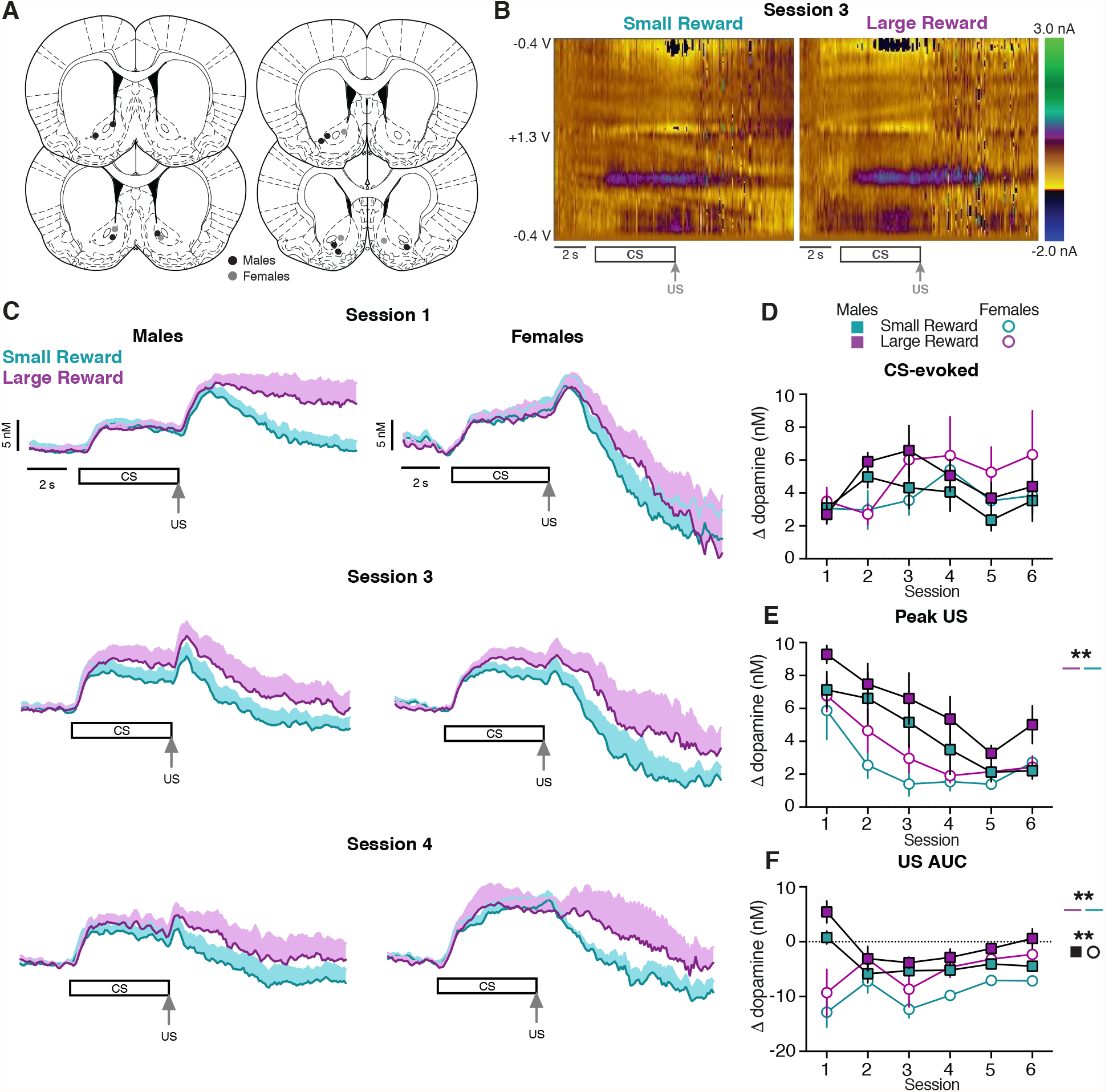
Dopamine release in the NAc during early training sessions. (A) Location of voltammetry electrodes in males (black) and females (gray). (B) Representative two-dimensional pseudocolor plots of the resulting current from voltage sweeps (y axis) as a function of time (x axis) of voltammetry recordings in the NAc. (C) Average dopamine signals across training sessions in males (left) and females (right). (D) Average CS-evoked dopamine release across sessions. (E) Average Peak US-evoked dopamine release across sessions. (F) Average US AUC-evoked dopamine release across sessions. * * *p* < 0.01.

To determine if reward-evoked dopamine release scaled with reward size, we quantified the maximum dopamine response during the 3 s after the reward was delivered relative to 0.5 s prior to the US delivery (peak US). Both male and female rats display a higher peak US dopamine response during Large Reward trials compared to Small Reward trials (three-way mixed-effects analysis; reward size effect: *F*_(1, 12)_ = 17.40, *p =* 0.001; **Fig. 3C, E**). We additionally analyzed the area under the curve for the average dopamine response during the 9 s after the US delivery relative to 0.5 s at the end of the CS (US AUC). Dopamine levels during this post-US period were higher following delivery of the Large Reward compared to the Small Reward (three-way mixed-effects analysis; reward size effect: *F*_(1, 12)_ = 17.98, *p* = 0.001; **Fig. 3F**). Furthermore, female rats displayed a lower US AUC dopamine response compared to male rats (three-way mixed-effects analysis; sex effect: *F*_(1, 30)_ = 7.91, *p* = 0.009; **Fig. 3F**). This difference in dopamine levels between trial types was also evident when examining the dopamine response during the Early and Late post-US epochs (**Supplementary Fig. 3**).

We next examined if the lower post-US dopamine levels in female rats could be explained by a difference in dopamine clearance. To address this the data were normalized to the peak US dopamine response and fit to a single-phase decay curve [32]. This analysis was performed on the first training session when there was a robust US dopamine response and only for the Small Reward trials to minimize the potential influence of multiple reward deliveries on the dynamics of the dopamine response. Females exhibited a decreased plateau (i.e., lower dopamine levels) compared to males (unpaired t-test, *t*_(9)_ = 2.65, *p =* 0.03; **Supplementary Fig. 4**). However, there was no difference in the tau between male and female rats (unpaired t-test, *t*_(9)_ = 0.92, *p =* 0.38; **Supplementary Fig. 4**), which indicates that the rate of the decay of the US-evoked dopamine response is not influenced by sex. Collectively, these results suggest that in contrast to CS-evoked dopamine release, US-evoked dopamine release encodes differences in reward size throughout the post-US period. Additionally, female rats exhibited a smaller US-evoked dopamine response relative to male rats.

CS-evoked dopamine release did not convey differences in reward size during early training sessions (**Fig. 4**). However, many studies demonstrate that the dopamine response to cues can convey differences in reward value in well-trained animals [3-8]. To determine if differences in CS-evoked dopamine emerge with further training, a subset of rats underwent three additional training sessions. In contrast to the first six training sessions, CS-evoked dopamine release signals differences in reward size in the following three training sessions (two-way mixed-effects analysis; reward size effect: *F*_(1, 10)_ = 5.78, *p* = 0.04; **Fig. 4A-B**). The peak US dopamine response did not differ by trial type in later sessions (two-way mixed-effects analysis; reward size effect: *F*_(1, 10)_ = 2.83, *p* = 0.12; **Fig. 4C**). However, the US AUC dopamine response remained higher following delivery of the Large Reward compared to the Small Reward (three-way mixed-effects analysis; reward size effect: *F*_(1, 10)_ = 24.54, *p* = 0.0006; **Fig. 4D**), and lower in female rats compared to male rats (three-way mixed-effects analysis; sex effect: *F*_(1, 10)_ = 8.76, *p* = 0.01; **Fig. 4D**), which is consistent with the findings from the first six training sessions. Furthermore, significant sex differences in conditioned responding and the post-US head entries were also evident during these later training sessions (**Supplementary Fig. 5**).

**Figure 4.**
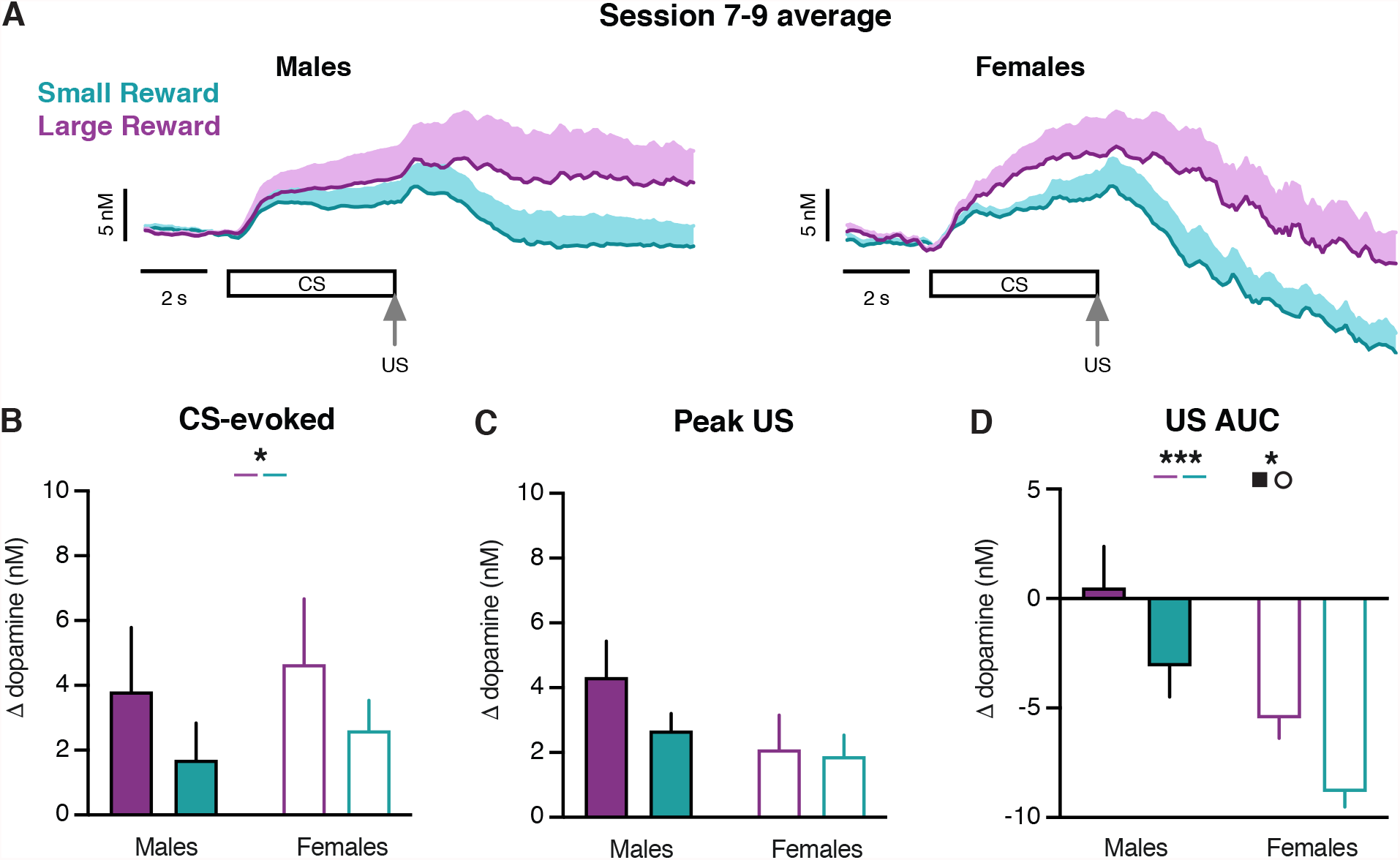
Dopamine release in the NAc during late training sessions. (A) Average of sessions 7-9 dopamine signals in males (left) and females (right). (B) Average CS-evoked dopamine release. (C) Average Peak US-evoked dopamine release. (D) Average US AUC-evoked dopamine release. * *p* < 0.05, * * * *p* < 0.001.

Prior studies have linked CS- and US-evoked dopamine release to conditioned responding [8,9,29,36-40]. Here, we utilized a repeated measures correlation analysis to determine how conditioned responding relates to dopamine transmission across all subjects and training sessions. While CS-evoked dopamine release was not correlated with conditioned responding (repeated measures correlation; conditioned responding: r_rm_ = −0.04, *p* = 0.60; latency: r_rm_ = 0.06, *p* = 0.40; **Fig. 5A**), there was an inverse relationship between conditioned responding and the peak US dopamine response (repeated measures correlation; r_rm_ = −0.15, *p* = 0.04; **Fig. 5B**). Furthermore, the number of head entries occurring in the 9 s following reward delivery was related with the US AUC dopamine response (repeated measures correlation; r_rm_ = 0.17, *p* = 0.02; **Supplementary Table**). These results highlight that behavioral responding during early Pavlovian learning is linked to US-evoked dopamine levels and unrelated to CS-evoked dopamine levels.

**Figure 5.**
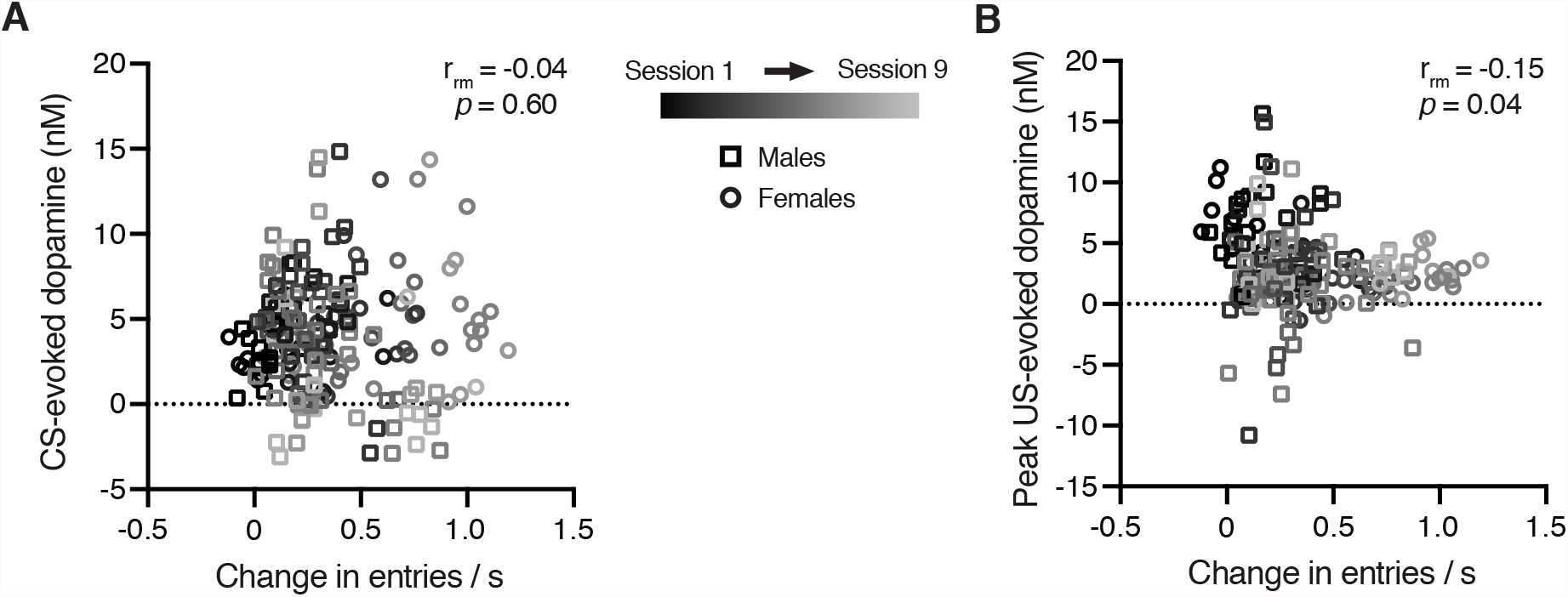
Relationship between dopamine and behavioral responding. (A) Relationship between CS-evoked dopamine release and conditioned responding. (B) Relationship between peak US-evoked dopamine release and conditioned responding.

## Discussion

The dopamine response to cues can signal differences in value-related information in well-trained animals [3-8]. For example, cue-evoked dopamine release conveys differences in the reward rate (i.e., the time elapsed since the previous reward delivery) after extensive training [8]. However, cue-evoked dopamine release does not signal differences in reward rate during the first six Pavlovian training sessions [9]. Our current results extend on these findings and demonstrate that during the first six training sessions, cue-evoked dopamine release did not signal differences in reward size. However with further training, we find reward size is encoded by the dopamine response to cues. Together this suggests that cue-evoked dopamine signals differences in reward value through a multistep process. First, cue-evoked dopamine signals an upcoming reward independent of value. Additional training is then required for cue-evoked dopamine release to encode differences in reward value.

Prior studies have identified sex differences in behavioral responding, as females display elevated motor activity compared to males in locomotor and anxiety-like assays [10-12]. Furthermore, sex differences have been observed across a variety of dopamine-dependent behaviors [13-15]. For example, female subjects display a faster acquisition rate and elevated responding during drug self-administration [16,17]. Females also exhibit higher levels of conditioned freezing relative to males during fear conditioning [18]. Sex differences have additionally been identified in Pavlovian conditioning tasks employing food rewards [19]. Specifically, female rats exhibited greater levels of sign-tracking behavior (e.g., physical interactions with a lever cue) but there were no sex differences in goal-tracking behavior (e.g., head entries to the food receptacle) [19]. By utilizing audio cues in our Pavlovian task, animals were not able to engage in standard sign-tracking behaviors. Regardless, we identified prominent sex differences in this Pavlovian task as females exhibited a higher level of goal-tracking compared to males. In addition to behavioral responses occurring during the cue, we analyzed the post-US head entries into the food port. Both males and females performed more head entries following the delivery of the large reward option. Furthermore, female rats performed more head entries compared to males throughout the post-US period. These findings illustrate previously unappreciated sex differences in behavioral responding following the delivery of rewards. We note that the sex differences observed across some of our behavioral metrics (latency and post-US head entries) could be explained by higher levels of motor activity in females [10-12]. However, our measure of conditioned responding is normalized to underlining differences in motor activity as we calculate the change in the rate of the head entries during the 5 s CS relative to the rate of head entries during the preceding 5 s. As such the sex differences in conditioned responding in our task cannot be explained solely by increased activity.

We identified stimulus-specific sex differences in dopamine release using fast-scan cyclic voltammetry. Cue-evoked dopamine release did not differ between males and females. In contrast, the reward-evoked dopamine response was lower in females relative to males. Prior research has identified lower levels of dopamine transmission in ovariectomized females compared to castrated males using microdialysis [26,41,42]. However, one must exercise caution generalizing the findings from ovariectomized females to intact females, as there are no sex differences in basal dopamine levels as measured by no net flux microdialysis within intact subjects [26]. Furthermore, a meta-analysis of microdialysis research concludes no basal or drug-induced sex differences in striatal dopamine [43]. As such, it is unlikely that any potential basal differences in dopamine levels between sexes could account for the stimulus-specific sex differences in rapid dopamine transmission.

Increasing evidence suggests that the estrous stage may contribute to the observed sex differences in dopamine release [23,24,26,41,42,44-48]. For example, the burst firing rate of dopamine neurons in the ventral tegmental area is elevated during estrous compared to other stages of the cycle in females, as well as compared to males [44,47]. Additional research finds that females elicit higher striatal dopamine release during estrous in response to electrical stimulation and cocaine, as measured using fast-scan cyclic voltammetry in anesthetized subjects [47]. We did not monitor the stages of the estrous cycle in the current study, so we cannot assess if these sex differences in reward-evoked dopamine release are due to cycling hormones. However, the observed sex differences were selective to the reward delivery (and not the cue) and were observed across sessions. Taken together, this evidence suggests the sex differences in dopamine transmission are not mediated by the estrous cycle and could instead reflect intrinsic differences between males and females. Additionally, we found no difference in the rate of decay of reward-evoked dopamine release between males and females, which suggests the observed sex difference is likely not due to differences in dopamine clearance. Future studies are needed to identify the source of these sex differences with in vivo dopamine transmission, which may result from anatomical and/or functional differences in the afferent input conveying reward-related information. Regardless, the lower US-evoked dopamine response in females could account for the results from human studies where females exhibit a diminished sensitivity to rewards relative to males [49].

The magnitude of the dopamine response to the cue presentation and reward delivery have been linked to behavioral outcomes in a variety of Pavlovian conditioning tasks [8,9,29,36-40]. We observed an inverse relationship between conditioned responding and reward-evoked dopamine release, which parallels our prior research and is consistent with the findings from studies employing optogenetic manipulations of the dopamine system [9,29,39]. However, our findings indicate conditioned responding does not differ by reward size and is not related to cue-evoked dopamine release within the first nine training sessions. In animals with extensive training in a similar Pavlovian task, an update in cue-evoked dopamine release can elicit differences in conditioned responding [8]. Taken together, these studies indicate the initial emergence of conditioned responding is linked to the decrease in reward-evoked dopamine release, whereas updates to cue-evoked dopamine release in well-trained animals leads to a corresponding update in conditioned responding.

## Acknowledgements

Special thanks to Dr. Claire Stelly for identifying and coding the repeated measures correlation analysis.

## Author Contributions

MJL and MID performed the experiments and analyzed the data. MJL and MJW designed the experiments and wrote the manuscript.

