## Supplementary Data for "Sex differences in behavioral responding and dopamine release during Pavlovian learning"

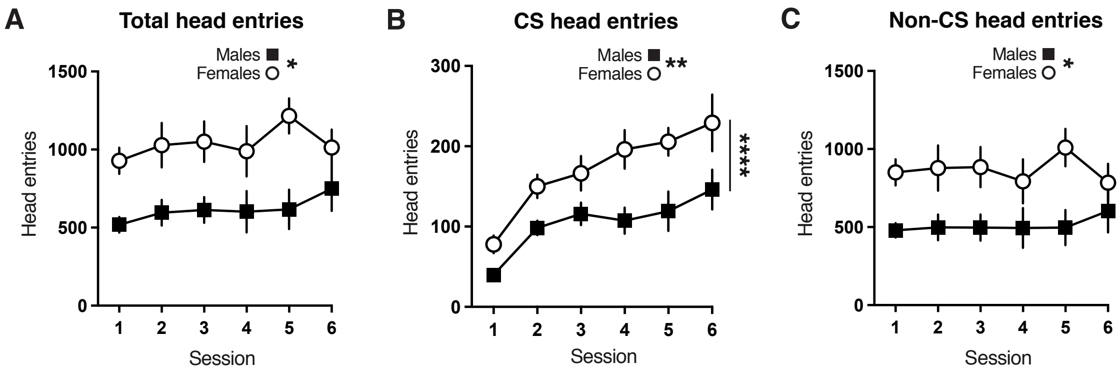


**Supplementary Figure 1.** Number of head entries across sessions. (A) Total head entries across sessions in males (black square) and females (open circle) (two-way mixed-effects analysis; session effect: F(5, 55) = 1.98, p = 0.10; sex effect: F(1, 11) = 8.38, p = 0.02; interaction effect: F(5, 55) = 1.25, p = 0.30). (B) CS head entries across sessions (two-way mixed-effects analysis; session effect: F(2.25, 26.69) = 15.34, p < 0.0001; sex effect: F(1, 11) = 12.10, p = 0.005; interaction effect: F(5, 55) = 0.97, p = 0.44). (C) Non-CS head entries across sessions (two-way mixed-effects analysis; session effect: F(2.47, 27.11) = 0.73, p = 0.52; sex effect: F(1, 11) = 6.12, p = 0.03; interaction effect: F(5, 55) = 1.59, p = 0.18). * $p < 0.05$, ** $p < 0.01$.

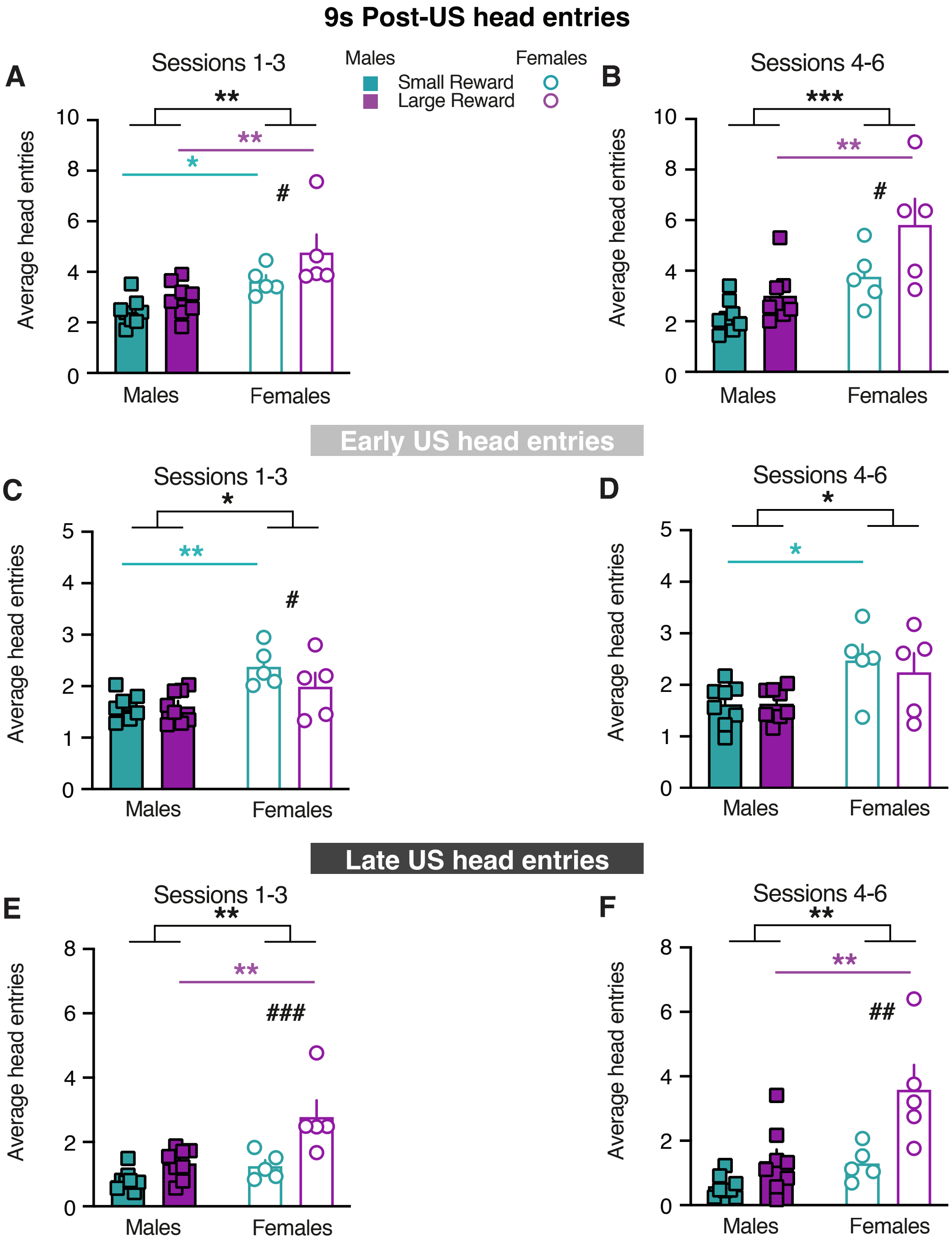


**Supplemental Figure 2.** Sex differences in behavioral responding during US presentation. (A) Head entries during the 9 s post-reward window averaged across the first three sessions of training (two-way mixed-effects analysis; reward size effect: $F(1, 11) = 15.03$, $p = 0.003$; sex effect: $F(1, 11) = 11.27$, $p = 0.006$; interaction effect: $F(1, 11) = 2.17$, $p = 0.17$; post hoc Sidak's test – sex; small reward: $t(22) = 2.41$, $p < 0.05$; large reward: $t(22) = 3.67$, $p = 0.003$; post hoc Sidak's test –

reward size; males: t(11) = 1.94, p = 0.15; females: t(11) = 3.41, p = 0.01). (B) Head entries during the 9 s post-reward window averaged across the latter three sessions of training (two-way mixed-effects analysis; reward size effect: F(1, 22) = 7.48, p = 0.01; sex effect: F(1, 22) = 17.45, p = 0.0004; interaction effect: F(1, 22) = 1.44, p = 0.24; post hoc Sidak's test – sex; small reward: t(22) = 2.11, p = 0.09; large reward: t(22) = 3.80, p = 0.002; post hoc Sidak's test – reward size; males: t(22) = 1.24, p = 0.41; females: t(22) = 2.51, p = 0.04). (C) Head entries during the Early US averaged across the first three sessions of training (two-way mixed-effects analysis; reward size effect: F(1, 11) = 5.47, p = 0.04; sex effect: F(1, 11) = 8.45, p = 0.01; interaction effect: F(1, 11) = 5.29, p = 0.04; post hoc Sidak's test – sex; small reward: t(22) = 3.57, p = 0.003; large reward: t(22) = 1.78, p = 0.17; post hoc Sidak's test – reward size; males: t(11) = 0.03, p = 0.99; females: t(11) = 2.96, p = 0.03). (D) Head entries during the Early US averaged across the latter three sessions of training (two-way mixed-effects analysis; reward size effect: F(1, 11) = 0.27, p = 0.61; sex effect: F(1, 11) = 9.67, p = 0.01; interaction effect: F(1, 11) = 0.35, p = 0.57; post hoc Sidak's test – sex; small reward: t(22) = 2.72, p = 0.02; large reward: t(22) = 1.94, p = 0.13; post hoc Sidak's test – reward size; males: t(11) = 0.71, p = 0.99; females: t(11) = 0.71, p = 0.74). (E) Head entries during the Late US averaged across the first three sessions of training (two-way mixed-effects analysis; reward size effect: F(1, 11) = 27.85, p = 0.0003; sex effect: F(1, 11) = 10.08, p = 0.009; interaction effect: F(1, 11) = 6.86, p = 0.02; post hoc Sidak's test – sex; small reward: t(22) = 1.22, p = 0.41; large reward: t(22) = 4.09, p = 0.001; post hoc Sidak's test – reward size; males: t(11) = 2.14, p = 0.11; females: t(11) = 5.03, p = 0.0008). (F) Head entries during the Late US averaged across the latter three sessions of training (two-way mixed-effects analysis; reward size effect: F(1, 11) = 17.75, p = 0.002; sex effect: F(1, 11) = 11.61, p = 0.006; interaction effect: F(1, 11) = 4.22, p = 0.06; post hoc Sidak's test – sex; small reward: t(22) = 1.26, p = 0.40; large reward: t(22) = 3.92, p = 0.002; post hoc Sidak's test – reward size; males: t(11) = 1.74, p = 0.21; females: t(11) = 3.99, p = 0.004). * denotes main effect of sex or post-hoc effect of sex, * $p < 0.05$, ** $p < 0.01$, *** $p < 0.001$. # denotes post-hoc effect of reward size, ## $p < 0.01$, ### $p < 0.001$.

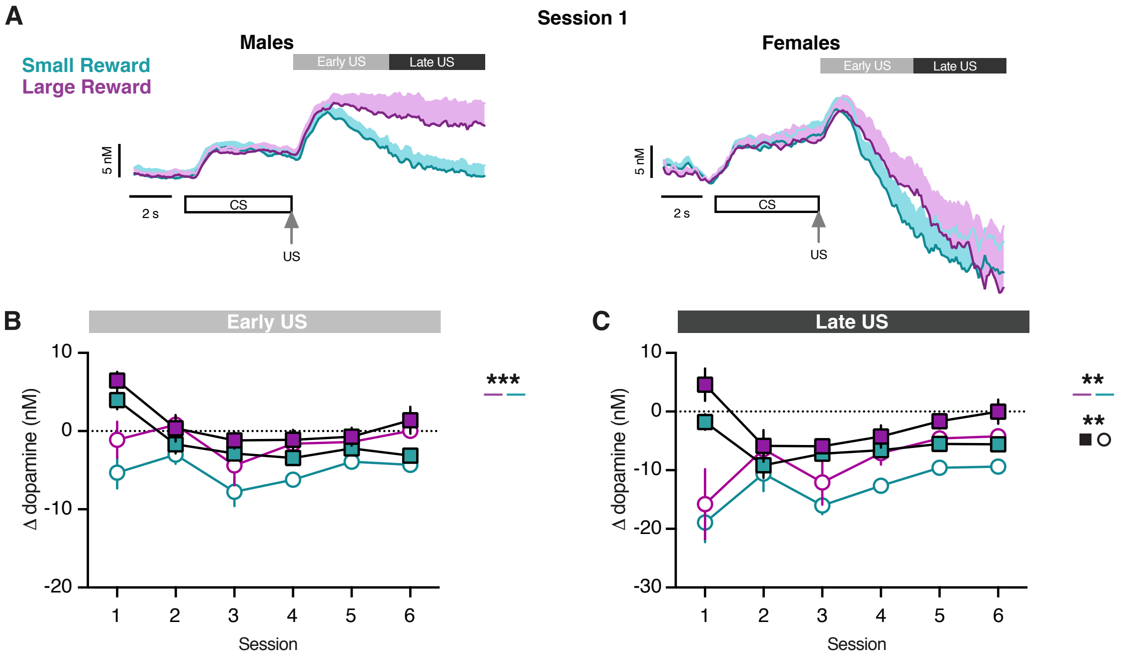


**Supplemental Figure 3.** Post-US dopamine response in Early and Late epochs. (A) Average dopamine signals in first session in males (left) and females (right) depicting Early and Late epochs. (B) Average Early US-evoked dopamine release across sessions (three-way mixed-effects analysis; session effect: F(2.77, 33.19) = 8.43, p = 0.0004; sex effect: F(1, 30) = 3.68, p = 0.06; reward size effect: F(1, 12) = 19.10, p = 0.0009; session x sex effect: F(5, 30) = 4.64, p = 0.003; session x reward size effect: F(2.38, 14.28) = 1.18, p = 0.34; sex x reward size effect: F(1, 30) = 1.01, p = 0.32; interaction effect: F(5, 30) = 1.22, p = 0.33). (C) Average Late US-evoked dopamine release across sessions (three-way mixed-effects analysis; session effect: F(1.80, 21.56) = 2.04, p = 0.16; sex effect: F(1, 30) = 9.49, p = 0.004; reward size effect: F(1, 12) = 15.32, p = 0.002; session x sex effect: F(5, 30) = 6.46, p = 0.0003; session x reward size effect: F(1.77, 10.62) = 0.58, p = 0.56; sex x reward size effect: F(1, 30) = 0.01, p = 0.93; interaction effect: F(5, 30) = 1.62, p = 0.18). ** $p < 0.01$, *** $p < 0.001$.

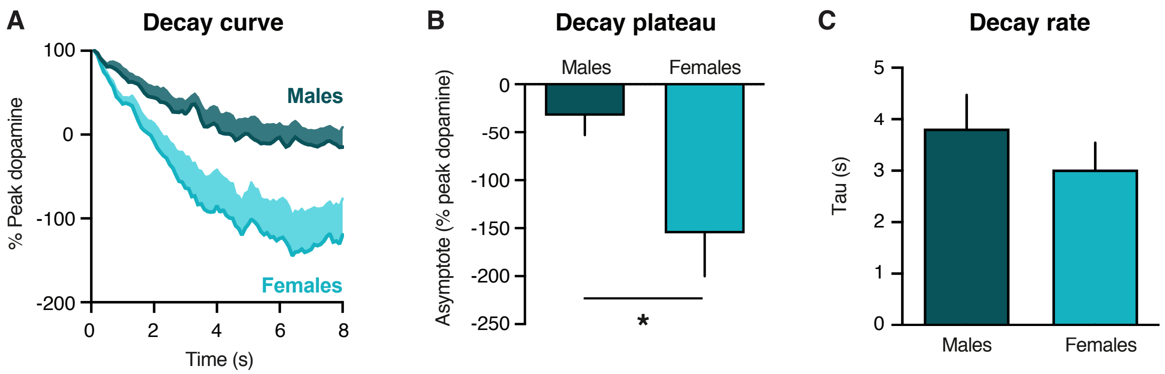


**Supplemental Figure 4.** Single-phase decay analysis on US dopamine response. (A) Average US-evoked dopamine signals normalized to the peak dopamine response in Small Reward trials during the first session in males and females. (B) Decay plateau (unpaired t-test; t(9) = 2.65, p = 0.03). (C) Decay rate (unpaired t-test; t(9) = 0.92, p = 0.38). * $p < 0.05$.

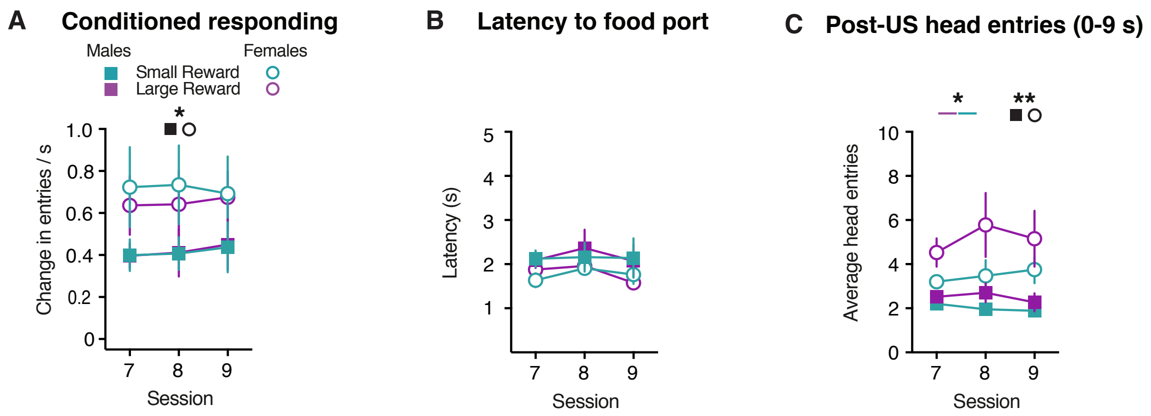


**Supplemental Figure 5.** Behavioral responding during sessions 7-9. (A) Conditioned responding (three-way mixed-effects analysis; session effect: F(1.99, 21.95) = 0.42, p = 0.66; sex effect: F(1, 18) = 5.56, p = 0.03; reward size effect: F(1, 11) = 0.11 p = 0.74; session x sex effect: F(2, 18) = 0.33, p = 0.72; session x reward size effect: F(1.76, 15.87) = 0.23, p = 0.77; sex x reward size effect: F(1, 18) = 0.13, p = 0.72; interaction effect: F(2, 18) = 0.13, p = 0.88). (B) Latency to respond (three-way mixed-effects analysis; session effect: F(1.68, 18.46) = 1.00, p = 0.37; sex effect: F(1, 11) = 1.68, p = 0.84; reward size effect: F(1, 11) = 0.04 p = 0.84; session x sex effect: F(2, 18) = 0.07, p = 0.94; session x reward size effect: F(1.83, 16.43) = 1.61, p = 0.23; sex x reward size effect: F(1, 18) = 0.003, p = 0.95; interaction effect: F(2, 18) = 1.51, p = 0.25). (C) Post-US head entries (three-way mixed-effects analysis; session effect: F(1.52, 16.67) = 0.86, p = 0.41; sex effect: F(1, 18) = 14.62, p = 0.001; reward size effect: F(1, 11) = 5.82 p = 0.03; session x sex effect: F(2, 18) = 1.46, p = 0.26; session x reward size effect: F(1.25, 11.20) = 0.93, p = 0.38; sex x reward size effect: F(1, 18) = 1.93, p = 0.18; interaction effect: F(2, 18) = 0.17, p = 0.84). * $p < 0.05$, ** $p < 0.01$.

| **Table 1 - Figure 1** (n = 8 males, n = 5 females) | | | |
|---|---|---|---|
| **Panel B – Conditioned responding** | | | |
| Three-way mixed-effects model | Session<br>$F_{(2.26, 24.86)}$ = 14.01, ***p* < 0.0001** | Sex<br>$F_{(1, 55)}$ = 3.90, *p* = 0.05 | Reward size<br>$F_{(1, 11)}$ = 0.03, *p* = 0.86 |
| Session x Sex<br>$F_{(5, 55)}$ = 2.34, *p* = 0.05 | Session x Reward size<br>$F_{(1.91, 20.97)}$ = 0.54, *p* = 0.58 | Sex x Reward size<br>$F_{(1, 55)}$ = 0.18, *p* = 0.67 | Three-way interaction<br>$F_{(5, 55)}$ = 1.20, *p* = 0.32 |
| **Panel C – Conditioned responding: Sessions 1-3** | | | |
| Two-way mixed-effects model | Reward size<br>$F_{(1, 11)}$ = 0.25, *p* = 0.63 | Sex<br>$F_{(1, 11)}$ = 0.90, *p* = 0.36 | Two-way interaction<br>$F_{(1, 11)}$ = 0.05, *p* = 0.83 |
| **Panel D – Conditioned responding: Sessions 4-6** | | | |
| Two-way mixed-effects model | Reward size<br>$F_{(1, 11)}$ = 0.50, *p* = 0.49 | Sex<br>$F_{(1, 11)}$ = 5.11, ***p* < 0.05** | Two-way interaction<br>$F_{(1, 11)}$ = 0.54, *p* = 0.48 |
| **Panel E – Latency to respond** | | | |
| Three-way mixed-effects model | Session<br>$F_{(2.94, 32.33)}$ = 6.26, ***p* < 0.002** | Sex<br>$F_{(1, 55)}$ = 8.80, ***p* = 0.004** | Reward size<br>$F_{(1, 11)}$ = 0.51, *p* = 0.49 |
| Session x Sex<br>$F_{(5, 55)}$ = 0.60, *p* = 0.70 | Session x Reward size<br>$F_{(2.12, 23.32)}$ = 1.40, *p* = 0.27 | Sex x Reward size<br>$F_{(1, 55)}$ = 0.72, *p* = 0.40 | Three-way interaction<br>$F_{(5, 55)}$ = 2.40, ***p* < 0.05** |
| **Panel F – Latency to respond: Sessions 1-3** | | | |
| Two-way mixed-effects model | Reward size<br>$F_{(1, 11)}$ = 1.71, *p* = 0.22 | Sex<br>$F_{(1, 11)}$ = 14.56, ***p* = 0.003** | Two-way interaction<br>$F_{(1, 11)}$ = 1.25, *p* = 0.29 |
| **Panel F – Latency to respond: Sessions 4-6** | | | |
| Two-way mixed-effects model | Reward size<br>$F_{(1, 11)}$ = 0.01, *p* = 0.94 | Sex<br>$F_{(1, 11)}$ = 2.71, *p* = 0.13 | Two-way interaction<br>$F_{(1, 11)}$ = 0.15, *p* = 0.71 |

| Table 2 - Figure 2 | | | |
|---|---|---|---|
| **Panel B –Post US head entries** | | | |
| Three-way mixed-effects model | Session<br>$F_{(1.98, 21.81)} = 1.30$, $p = 0.29$ | Sex<br>$F_{(1, 55)} = 17.44$, $\boldsymbol{p = 0.0001}$ | Reward size<br>$F_{(1, 11)} = 10.15$, $\boldsymbol{p = 0.009}$ |
| Session x Sex<br>$F_{(5, 55)} = 1.79$, $p = 0.13$ | Session x Reward size<br>$F_{(2.52, 27.69)} = 1.62$, $p = 0.21$ | Sex x Reward size<br>$F_{(1, 55)} = 1.77$, $p = 0.19$ | Three-way interaction<br>$F_{(5, 55)} = 0.94$, $p = 0.46$ |
| **Panel C – Early US head entries** | | | |
| Three-way mixed-effects model | Session<br>$F_{(2.49, 27.40)} = 1.72$, $p = 0.19$ | Sex<br>$F_{(1, 55)} = 13.60$, $\boldsymbol{p = 0.0005}$ | Reward size<br>$F_{(1, 11)} = 1.45$, $p = 0.25$ |
| Session x Sex<br>$F_{(5, 55)} = 0.99$, $p = 0.43$ | Session x Reward size<br>$F_{(3.02, 33.24)} = 0.68$, $p = 0.57$ | Sex x Reward size<br>$F_{(1, 55)} = 1.54$, $p = 0.22$ | Three-way interaction<br>$F_{(5, 55)} = 0.93$, $p = 0.47$ |
| **Panel D – Late US head entries** | | | |
| Three-way mixed-effects model | Session<br>$F_{(1.65, 18.10)} = 0.67$, $p = 0.50$ | Sex<br>$F_{(1, 55)} = 12.20$, $\boldsymbol{p = 0.001}$ | Reward size<br>$F_{(1, 11)} = 24.65$, $\boldsymbol{p = 0.0004}$ |
| Session x Sex<br>$F_{(5, 55)} = 1.83$, $p = 0.12$ | Session x Reward size<br>$F_{(2.01, 22.15)} = 2.92$, $p = 0.07$ | Sex x Reward size<br>$F_{(1, 55)} = 5.94$, $\boldsymbol{p = 0.02}$ | Three-way interaction<br>$F_{(5, 55)} = 1.50$, $p = 0.21$ |

| Table 3 - Figure 3 (males: n = 9 electrodes; females: n = 5 electrodes) | | | |
|---|---|---|---|
| **Panel D – CS-evoked dopamine release** | | | |
| Three-way mixed-effects model | Session<br>$F_{(1.97, 23.64)} = 3.22$, $p = 0.06$ | Sex<br>$F_{(1, 30)} = 0.07$, $p = 0.80$ | Reward size<br>$F_{(1, 12)} = 3.54$, $p = 0.09$ |
| Session x Sex<br>$F_{(5, 30)} = 2.48$, $p = 0.05$ | Session x Reward size<br>$F_{(2.78, 16.70)} = 2.78$, $p = 0.08$ | Sex x Reward size<br>$F_{(1, 30)} = 0.19$, $p = 0.67$ | Three-way interaction<br>$F_{(5, 30)} = 0.63$, $p = 0.68$ |
| **Panel E – Peak US-evoked dopamine release** | | | |
| Three-way mixed-effects model | Session<br>$F_{(2.43, 29.14)} = 13.18$, $\mathbf{p < 0.0001}$ | Sex<br>$F_{(1, 30)} = 3.14$, $p = 0.09$ | Reward size<br>$F_{(1, 12)} = 17.40$, $\mathbf{p = 0.001}$ |
| Session x Sex<br>$F_{(5, 30)} = 0.47$, $p = 0.80$ | Session x Reward size<br>$F_{(1.36, 8.14)} = 0.27$, $p = 0.69$ | Sex x Reward size<br>$F_{(1, 30)} = 1.97$, $p = 0.17$ | Three-way interaction<br>$F_{(5, 30)} = 1.73$, $p = 0.16$ |
| **Panel F – AUC US-evoked dopamine release** | | | |
| Three-way mixed-effects model | Session<br>$F_{(2.06, 24.72)} = 2.54$, $p = 0.10$ | Sex<br>$F_{(1, 30)} = 7.91$, $\mathbf{p = 0.009}$ | Reward size<br>$F_{(1, 12)} = 17.98$, $\mathbf{p = 0.001}$ |
| Session x Sex<br>$F_{(5, 30)} = 6.56$, $\mathbf{p = 0.0003}$ | Session x Reward size<br>$F_{(1.96, 11.74)} = 0.65$, $p = 0.54$ | Sex x Reward size<br>$F_{(1, 30)} = 0.18$, $p = 0.67$ | Three-way interaction<br>$F_{(5, 30)} = 1.36$, $p = 0.27$ |

| Table 4 - Figure 4 | | | |
|---|---|---|---|
| **Panel B – CS-evoked dopamine: Sessions 7-9** | | | |
| Two-way mixed-effects model | Reward size<br>$F_{(1, 10)} = 5.78$, $\boldsymbol{p = 0.04}$ | Sex<br>$F_{(1, 10)} = 0.16$, $p = 0.70$ | Two-way interaction<br>$F_{(1, 10)} = 0.001$, $p = 0.97$ |
| **Panel C – Peak US dopamine: Sessions 7-9** | | | |
| Two-way mixed-effects model | Reward size<br>$F_{(1, 10)} = 2.83$, $p = 0.12$ | Sex<br>$F_{(1, 10)} = 1.74$, $p = 0.22$ | Two-way interaction<br>$F_{(1, 10)} = 1.68$, $p = 0.22$ |
| **Panel D – AUC US dopamine: Sessions 7-9** | | | |
| Two-way mixed-effects model | Reward size<br>$F_{(1, 10)} = 24.54$, $\boldsymbol{p = 0.0006}$ | Sex<br>$F_{(1, 10)} = 8.76$, $\boldsymbol{p = 0.01}$ | Two-way interaction<br>$F_{(1, 10)} = 0.03$, $p = 0.87$ |

| Table 5: Figure 5 | |
|---|---|
| **Sessions 1-9 Repeated measures correlation** | |
| **Panel A – CS dopamine vs conditioned responding** | $r_{rm}$ = -0.04, *p* = 0.60 |
| **Panel B – peak US dopamine vs conditioned responding** | $r_{rm}$ = -0.15, ***p* = 0.04** |
| **Additional analyses** | |
| **CS dopamine vs latency to respond** | $r_{rm}$ = 0.06, *p* = 0.40 |
| **AUC US dopamine vs 9s post-US HE** | $r_{rm}$ = 0.17, ***p* = 0.02** |
| **AUC US dopamine vs conditioned responding** | $r_{rm}$ = -0.02, *p* = 0.82 |
